# Molecular signatures of genetically defined jugular nociceptors and their responses to allergic airway inflammation

**DOI:** 10.64898/2026.09.24.753851

**Authors:** Yiqi Wang, Katy Lawson, Ryan Stewart, Shizhen Tang, Shuang Wu, Margaret Y Chang, Vishnu V Yella, Haylie L. Ramsey, Jingjing Yang, Liang Han

## Abstract

**Rationale:** Vagal sensory neurons are molecularly and functionally diverse, and they play critical roles in visceral sensation and homeostasis. The vagal system is anatomically divided into jugular and nodose ganglia, which exhibit distinct peripheral innervation patterns and physiological functions. Although recent transcriptomic studies have characterized nodose neuronal subtypes in detail, the molecular profiles of jugular sensory neurons remain poorly defined.

**Objectives:** To define the molecular profiles of genetically identified jugular sensory neurons and determine the transcriptional responses of jugular and nodose airway afferents to allergic airway inflammation.

**Methods:** We performed deep transcriptomic profiling of two genetically defined jugular nociceptor populations, MrgprC11⁺ and MrgprD⁺ neurons, and compared them with airway-innervating P2ry1⁺ nodose neurons and Phox2b⁺ nodose neurons. We further examined transcriptional changes of MrgprC11⁺ and P2ry1⁺ neurons following OVA-induced allergic airway inflammation.

**Measurements and Main Results:** Transcriptomic analyses revealed distinct molecular signatures of jugular nociceptive neurons and identified subtype-specific genes across diverse functional categories. Following OVA challenge, MrgprC11⁺ and P2ry1⁺ neurons exhibited strikingly divergent transcriptional responses. More than 400 genes that were differentially expressed in both neuronal populations were regulated in opposite directions. GO enrichment analysis suggested that MrgprC11⁺ jugular neurons adopted a metabolically restrained state with reduced translation and bioenergetic activities, whereas P2ry1⁺ nodose neurons exhibited metabolically active state with enhanced translation and bioenergetic activities.

**Conclusions:** These findings provide new insight into vagal sensory neuron diversity and reveal distinct transcriptional adaptations of jugular and nodose airway afferents during allergic inflammation.

## Introduction

Vagal sensory neurons detect diverse mechanical, chemical, and inflammatory stimuli arising from visceral organs and relay this interoceptive information to the central nervous system. By monitoring the physiological state of the respiratory, gastrointestinal, and cardiovascular systems, these neurons regulate autonomic reflexes essential for maintaining physiological homeostasis (1–3). Vagal sensory neurons arise from two anatomically and developmentally distinct ganglia, the placode-derived nodose ganglion and the neural crest-derived jugular ganglion, with nodose neurons accounting for approximately 80% and jugular neurons approximately 20% of the vagal sensory neuron population (4, 5). Recent advances in transcriptomic profiling, anatomical mapping, and functional studies revealed that vagal sensory neurons comprise numerous molecularly and functionally distinct subtypes (5–14), many characterized by unique peripheral innervation patterns, response properties, and physiological functions. In particular, multiple nodose neuron subtypes have been molecularly defined and linked to distinct visceral organs, where they mediate specialized physiological functions, including respiratory regulation, gastrointestinal stretch detection and nutrient sensing, and cardiovascular reflexes (5, 8, 11, 12, 15–21). These studies have transformed our understanding of vagal sensory circuits by linking molecular identity to organ-specific sensory functions.

However, comprehensive transcriptomic characterization of jugular sensory neurons has remained technically challenging because jugular neurons constitute only a small fraction of the vagal ganglion and exhibit poor survival during tissue dissociation. Consequently, most transcriptomic studies of vagal sensory neurons have provided little or no information on jugular neuron populations (5, 6, 9, 11, 12, 14). Although several existing transcriptomic datasets revealed information of jugular neuronal subtypes (7, 8, 10, 13), providing an important molecular framework for studying jugular sensory neurons, the absence of a comprehensive molecular characterization of genetically defined jugular neuron subtypes has become a major barrier to understanding their physiological functions in visceral interoception and homeostatic regulation.

The Mas-related G protein-coupled receptor (Mrgpr) family comprises more than 20 functional receptors that are classified into four subfamilies: MrgprA, MrgprB, MrgprC, and MrgprD (22). Initially identified as sensory neuron-specific receptors in dorsal root ganglia, Mrgprs have been shown to play important roles in somatosensation, including pain, itch, and touch (23–26). Our recent studies discovered that Mrgpr family members are selectively expressed in the jugular neurons, but not nodose neurons. Expression of MrgprC11 and MrgprD define two distinct subsets of jugular nociceptor populations (27). Functional studies further demonstrated that MrgprC11⁺ jugular neurons densely innervate the airway and regulate bronchoconstriction and airway hyperresponsiveness (27, 28), whereas MrgprD⁺ jugular neurons do not innervate the lung and their peripheral targets and physiological functions remain unknown (29). The availability of the genetic tools labeling MrgprC11^+^ and MrgprD^+^ neurons (27, 30) provide the unique opportunity for molecular profiling of jugular nociceptive populations.

Sensory neurons exhibit remarkable molecular plasticity in response to peripheral inflammation. Tissue injury and inflammatory mediators can remodel neuronal transcriptional programs, alter neuronal excitability, and ultimately reshape sensory functions. Indeed, recent studies have shown that allergic airway inflammation induces significant transcriptional changes of airway sensory neurons, while inflammatory cytokines directly regulate gene expression (31–33). For instance, our previous study showed that MrgprC11⁺ jugular sensory neurons are sensitized by allergic inflammatory cytokines, including IL-4 and oncostatin M (27). Airway constriction induced by the activation of MrgprC11^+^ jugular neurons was also enhanced in the asthmatic mice, suggesting that allergic airway inflammation sensitized MrgprC11^+^ jugular neurons (27). However, how allergic lung inflammation alters the transcriptome of MrgprC11^+^ neurons and subsequently leads to cellular dysfunction remains unclear. Moreover, whether distinct subsets of airway-innervating neurons undergo similar or distinct transcriptional changes during allergic airway inflammation is not clear.

Here, combining Cre-dependent genetic labeling, fluorescence-activated cell sorting, and ultra-sensitive, low-input SMART-seq strategy, we performed transcriptomic analysis of MrgprC11⁺ and MrgprD⁺ jugular nociceptors. We compared these populations with Phox2B⁺ nodose neurons and P2ry1⁺ airway-innervating nodose A-fibers to identify jugular-enriched molecular signatures. Moreover, to define how vagal sensory neuron respond to allergic lung inflammation, we examined transcriptomic changes of MrgprC11^+^ jugular nociceptors and P2ry1⁺ airway-innervating nodose A-fibers under naïve and allergic airway inflammation conditions. Our results show that allergic airway inflammation does not induce a uniform response across vagal sensory neurons; instead, MrgprC11⁺ and P2ry1⁺ neurons undergo fundamentally distinct and opposite transcriptional adaptations to the same inflammatory stress.

## Results

### Four vagal sensory neuronal populations for RNA sequencing analysis: MrgprC11^+^, MrgprD^+^, P2ry1^+^, and Phox2B^+^ neurons

Our recent work demonstrated that MrgprC11 and MrgprD mark two molecularly distinct populations of small-diameter C-fiber neurons within the jugular ganglion (Figure 1B), comprising 20.3% and 37.3% of jugular neurons, respectively (27). Functional studies further established that MrgprC11⁺ jugular sensory neurons play a critical role in regulating bronchoconstriction and airway hyperresponsiveness (27, 28). In contrast, the physiological function of MrgprD^+^ jugular neurons remains unknown. Using *MrgprC11^CreER^* and *MrgprD^CreER^*mouse lines (27, 30), we genetically labeled these neuronal populations for fluorescence-activated cell sorting (FACS) followed by RNA sequencing (Table 1, Figure 1A).

**Figure 1.**
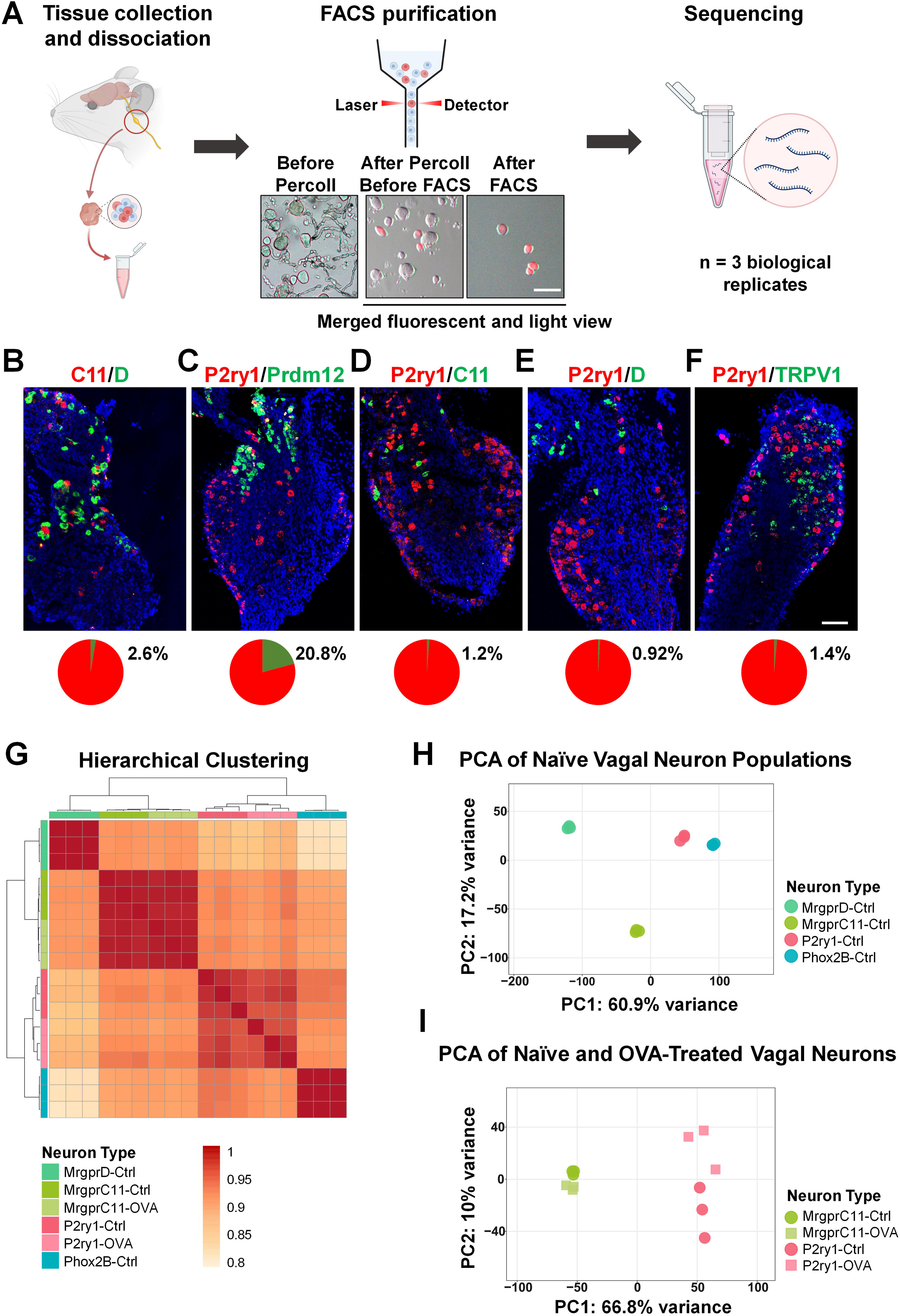
Transcriptomic profiling of four vagal sensory neuronal populations: MrgprC11⁺, MrgprD⁺, airway-innervating P2ry1⁺, and Phox2b^+^ neurons. (A) Schematic of the FACS-based neuronal isolation and RNA sequencing workflow. Vagal ganglia from *MrgprC11^tdTomato^, MrgprD^tdTomato^, P2ry1^tdTomato^, or Phox2b^tdTomato^*mice were dissociated, enriched by Percoll gradient centrifugation, and sorted by FACS to isolate labeled neurons. Representative images in the middle show progressive enrichment before Percoll, after Percoll, and after FACS. (B-F). Top: RNAscope in situ hybridization detecting indicated markers. DAPI nuclear staining is shown in blue. Bottom: Quantification of marker overlap. Values indicate the percentage of neurons labeled by the green marker within the population labeled by the red marker. C11: MrgprC11, D: MrgprD, Scale bars: 100 µm. (G) Hierarchical clustering showing high replicate consistency within each neuronal population. (H) PCA of four naïve vagal sensory neuron populations. (I) PCA of naïve and OVA-treated MrgprC11⁺ and P2ry1⁺ neurons, showing treatment-associated transcriptomic shifts following allergic airway inflammation.

**Table 1.**
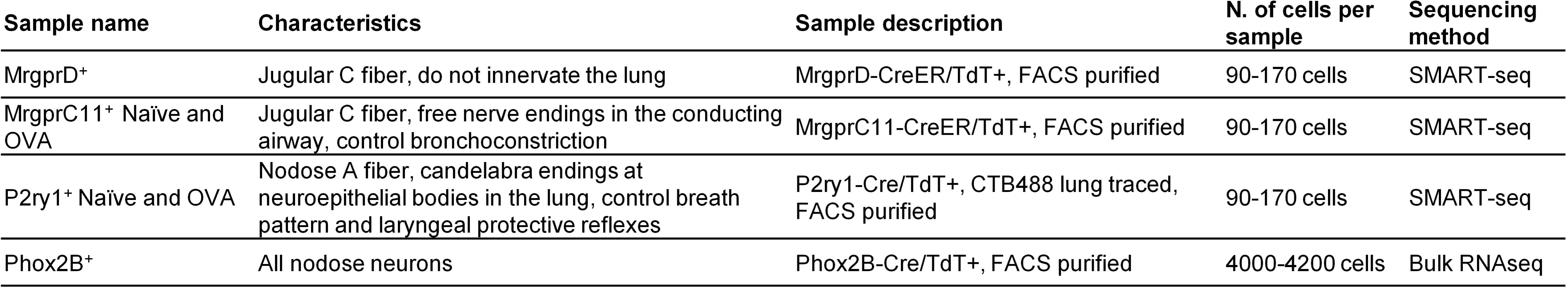
Summary of FACS-purified vagal sensory neuron populations used for transcriptomic profiling.

To understand how allergic inflammation in the lung changes the transcriptome of vagal sensory neurons, we performed RNAseq of MrgprC11⁺ jugular neurons and P2ry1^+^ airway-innervating neurons isolated from both naïve and asthmatic mice (Table 1). P2ry1^+^ vagal sensory nerves in the lung are predominantly nodose-derived myelinated A-fibers and constitute the majority of airway-innervating A fibers (8, 15, 34). They form stereotyped candelabra terminals at neuroepithelial bodies in the lung and regulate breathing patterns (15). More recent work further demonstrated that vagal P2ry1 neurons are required for protective responses to laryngeal water and acid challenge, establishing this population as a key component of airway protective reflexes (8). Notably, RNAscope *in situ* hybridization demonstrated that P2ry1⁺ neurons were molecularly distinct from MrgprC11⁺ and MrgprD⁺ jugular neurons. P2ry1 exhibited negligible co-expression with MrgprC11 (1.2%) and MrgprD (0.92%) (Figure 1D-E). Moreover, the majority of P2ry1⁺ neurons (79.2%) do not express the jugular marker Prdm12 (Figure 1C). Furthermore, P2ry1 expression showed minimal overlap with the nociceptive marker TRPV1 (1.4%), consistent with the fact that P2ry1 mainly labels myelinated A-fibers (Figure 1F). Because P2ry1⁺ neurons represent a molecularly distinct airway-innervating nodose A-fiber population, we included them as a nodose comparison group to MrgprC11⁺ jugular neurons for examining cell-type-specific responses to allergic lung inflammation.

Our analyses also included Phox2B⁺ nodose neurons as a reference population representing the overall nodose ganglion (7).

### Isolation of neurons and sequencing strategy

Phox2B⁺ neurons were isolated from *Phox2B^Cre^; ROSA26^tdTomato^* mice. To isolate airway-innervating P2ry1^+^ vagal sensory neurons, CTB-488 was delivered via intratracheal instillation into *P2ry1^Cre^; ROSA26^tdTomato^* mice. Cells were isolated by FACS based on tdTomato and CTB488 fluorescence. Due to the low abundance of MrgprC11⁺ and MrgprD⁺ jugular neurons, selective isolation of airway-innervating subsets was technically infeasible. Therefore, all tdTomato^+^ cells were collected from *MrgprC11^CreER^; ROSA26^tdTomato^* and *MrgprD^CreER^; ROSA26^tdTomato^*respectively (Table 1).

Because vagal sensory neuron subpopulations differ in abundance, we used an input-dependent sequencing strategy. Low-abundance MrgprC11⁺, MrgprD⁺, and P2ry1⁺ neurons (90-170 cells per sample) were profiled by SMART-seq, which was designed for low-input transcriptomic analysis (35). Phox2B⁺ neurons were included to establish a transcriptomic reference representative of the nodose ganglion in our analysis. Because each mouse nodose ganglion contains thousands of neurons spanning numerous molecularly distinct subtypes, we collected a large number of Phox2B⁺ neurons (4000-4200 cells per sample) to maximize representation of the *in vivo* nodose neuronal diversity. Therefore, the Phox2B⁺ population was profiled by bulk RNA sequencing rather than SMART-seq. All samples were sequenced on an Illumina HiSeq 2500 platform to an average depth of ∼32 million mapped reads per sample (range: 17.8-46.0 million), with an average of 20,426 genes (range: 16,433-24,046) detected per sample across all sequenced populations.

### Hierarchical clustering and principal components analysis

To compare transcriptomic relationships among vagal sensory neuron subtypes, we performed hierarchical clustering and principal component analysis (PCA) utilizing variance-stabilizing transformed (VST) to count matrices across all quantified genes (Figure 1G-I). Sample-to-sample correlation analysis demonstrated strong technical reproducibility and group homogeneity, with replicates within each distinct subtype forming tight, highly correlated clusters (*r* ≥ 0.97, with group means ranging from 0.971 to 0.996) (Figure 1G).

Principal component analysis of naïve vagal neuron populations revealed that PC1 explained 60.9% of the variance and clearly separated samples according to embryonic ganglion origin. The nodose populations, P2ry1-Ctrl and Phox2B-Ctrl, clustered together on the right side of PC1, whereas the jugular populations, MrgprD-Ctrl and MrgprC11-Ctrl, separated to the left. PC2, which explained 17.2% of the variance, further separated the two jugular populations, identifying MrgprD⁺ and MrgprC11⁺ neurons as transcriptomically distinct subpopulations (Figure 1H). Notably, under OVA-treated conditions, both MrgprC11⁺ and P2ry1⁺ populations clustered closely with their respective naïve controls in the focused PCA analysis (PC1: 66.8%; PC2: 10.0%), with only modest treatment-associated shifts along the secondary axis. This suggests that allergic airway inflammation changes the transcriptomic state of these neurons without changing their core cell-type identity (Figure 1I).

### Distinct expression of physiological sensors, transducers, and synaptic molecules in different subtypes of neurons

We first examine the expression of genes that are important for stimuli detection, sensory transduction, action potential generation, and neurotransmitter release in MrgprC11⁺, MrgprD⁺, and P2ry1⁺-airway innervating neurons (Figure 2, Table S1). The three populations showed expression patterns consistent with known nodose and jugular markers (Figure 2A) (4, 7, 12, 27). The established nodose markers *P2rx2, Htr3a,* and *Htr3b* were selectively expressed in P2ry1⁺ neurons, whereas *Mrgpr* family members were selectively expressed in MrgprC11⁺ and MrgprD⁺ jugular neurons. Neuropeptides *Tac1* and *Calca/Calcb* were also highly enriched in the jugular populations.

**Figure 2.**
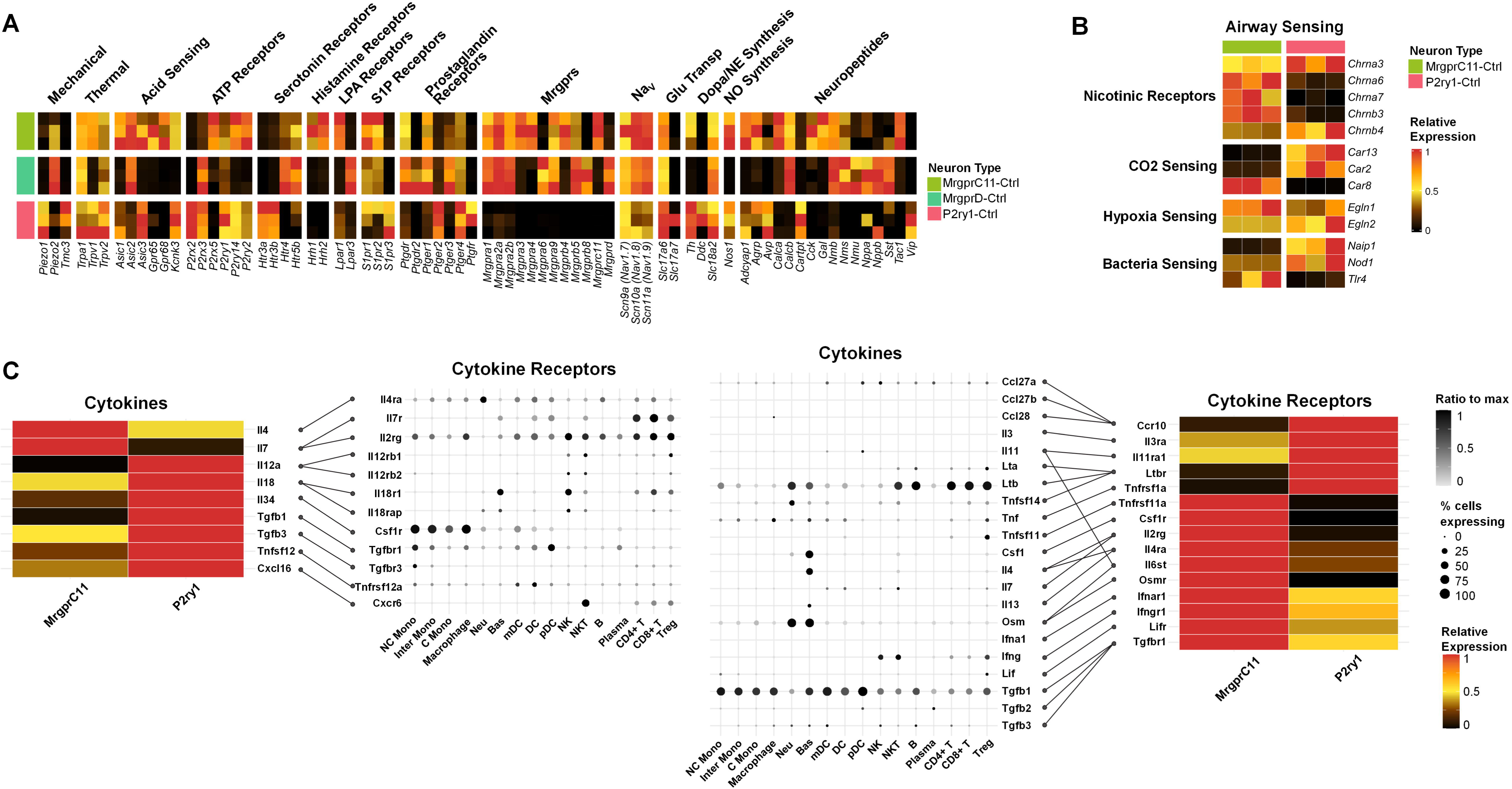
Functional gene expression across different neuronal populations. (A) Heatmap of relative expression (row-normalized, 0–1) for key sensory genes across MrgprC11⁺, MrgprD⁺, and P2ry1⁺ neuron populations. Color indicates expression relative to each gene’s maximum across the three populations. (B) Expression for key genes involved in airway sensing in MrgprC11^+^ and P2ry1^+^ neurons. The full gene lists for heatmap in A and B are provided in Table S1. (C) Potential bidirectional cytokine signaling between sensory neurons and lung immune cells. Heatmaps show differentially expressed cytokines and receptors between MrgprC11⁺ and P2ry1⁺ neurons, connected to their cognate receptors or ligands across immune cell types in the Tabula Muris Lung Atlas. Connecting lines indicate ligand-receptor pairs. Dot size represents the percentage of cells expressing each gene, and color indicates expression relative to the maximum for each gene. Abbreviations: NC Mono, non-classical monocyte; Inter Mono, intermediate monocyte; C Mono, classical monocyte; Neu, neutrophil; Bas, basophil; mDC, myeloid dendritic cell; DC, dendritic cell; pDC, plasmacytoid dendritic cell; NK, natural killer cell; NKT, natural killer T cell; B, B cell; Plasma, plasma cell; CD4⁺ T, CD4⁺ alpha-beta T cell; CD8⁺ T, CD8⁺ alpha-beta T cell; Treg, regulatory T cell. The full gene list is provided in Table S2.

Clear differences were also observed between the C-fiber populations (MrgprC11⁺/MrgprD⁺) and P2ry1⁺ A-fibers, largely resembling the molecular distinctions previously established between C-and A-fiber populations in the DRG. Sensory neuron-specific voltage-gated sodium channels exhibited fiber-type-associated expression patterns: *Scn9a* (NaV1.7) was highly expressed across both A-and C-fiber populations, whereas *Scn10a* (NaV1.8) and *Scn11a* (NaV1.9) were preferentially enriched in the jugular C-fiber populations. *Trpv2* expression was substantially higher in P2ry1⁺ neurons, consistent with its preferential expression in myelinated sensory neurons, whereas *Trpv1* was absent from MrgprD⁺ neurons, similar to the molecular profile of MrgprD⁺ DRG neurons (25, 36). *Trpa1* and *Trpv1* were detected in jugular C-fibers as well as P2ry1⁺ neurons due to the presence of a small unmyelinated P2ry1⁺ population. *Piezo1* and the putative mechanosensory gene *Tmc3* were more highly expressed in P2ry1⁺ neurons. Although *Piezo2* was expressed across all three populations, its expression was highest in MrgprD⁺ neurons. This mirrors the findings in DRG neurons, where Piezo2 is expressed in most of MrgprD⁺ neurons and its conditional deletion abolish mechanically activated currents in approximately half of these cells (37).

GPCRs play central roles in sensory neurons by detecting diverse chemical signals and regulating neuronal activity. Interestingly, many GPCR families, including serotonin receptors, histamine receptors, lysophosphatidic acid (LPA) receptors, Sphingosine 1-phosphate (S1P) receptors, and prostaglandin receptors, showed distinct expression patterns among the three neuronal populations. For instance, *Htr4* and *Htr5b* were enriched in jugular neurons. The two detected histamine receptors, *Hrh1* and *Hrh2*, were selectively enriched in MrgprC11⁺ neurons. LPA receptors were selectively expressed in jugular neurons, with *Lpar1* enriched in MrgprC11⁺ neurons and *Lpar3* in MrgprD⁺ neurons. In addition, the proton-sensing GPCRs *Gpr65* and *Gpr68* were selectively enriched in MrgprC11⁺ neurons, consistent with the role of these neurons as nociceptors with free nerve endings positioned in the conducting airways to detect chemical and inflammatory stimuli (27).

In summary, the three subsets of neurons exhibited distinct expressions of receptors, ion channels, and molecular components underlying sensory neuron function, suggesting that each population possesses specialized molecular machinery for detecting and transmitting sensory information.

### Distinct molecular programs for airway sensing and neuroimmune interactions

Sensory nerves in the airway are also equipped to detect a broad range of chemical and biological challenges, such as environmental chemicals and pathogen signals. We therefore examined genes associated with these sensory functions in two airway-innervating groups: MrgprC11⁺ and P2ry1⁺ neurons (Figure 2B, Table S1). Similarly, the two subtypes of neurons showed distinct expression patterns of nicotinic acetylcholine receptor subunits, carbonic anhydrases (*Car2, Car8, and Car13*) associated with CO₂/pH regulation, hypoxia-responsive enzymes (*Egln1 and Egln2*), and microbial pattern-recognition receptors (*Naip1, Nod1, and Tlr4*).

We next evaluated the capacity of MrgprC11⁺ and P2ry1⁺ neurons to participate in pulmonary neuroimmune signaling. We examined cytokine and cytokine receptor expressions in each neuronal population and connected them to their cognate receptors or ligands expressed by immune cells in the mouse lung cell atlas (Figure 2C, Table S2). Because comprehensive single-cell datasets of the inflamed lung remain limited, we focused on naïve sensory neurons and the available healthy lung cell atlas (38). Both neuron populations expressed multiple cytokines and cytokine receptors, similar to other airway-innervating sensory neuron populations (12), indicating broad roles of airway sensory neurons in neuroimmune signaling. However, their distinct repertoires suggest subtype-specific pathways for pulmonary neuroimmune communication.

For instance, *Lifr* and *Osmr*, which encode receptors for the IL-6 family cytokines LIF and OSM, respectively, were highly expressed in MrgprC11⁺ neurons (Figure 2C). Consistently, *Il6st* (gp130), the common signaling co-receptor for IL-6 family cytokines, was also highly enriched in MrgprC11⁺ neurons. LIF is produced primarily by non-classical monocytes and regulatory T cells, whereas OSM is predominantly associated with basophils and neutrophils, suggesting preferential signaling from these immune populations to MrgprC11⁺ jugular neurons. Indeed, our previous study suggested that OSM sensitizes MrgprC11⁺ jugular neurons during allergic airway inflammation and enhances airway hyperresponsiveness (27). Notably, most cytokines produced by basophils, including TGFβ1, OSM, IL-4, and CSF1, have corresponding receptors enriched in MrgprC11⁺ neurons, suggesting that basophils may represent an important immune cell partner for MrgprC11⁺ jugular neurons.

Conversely, MrgprC11⁺ neurons showed enriched expression of the cytokines *Il4* and *Il7* compared with P2ry1⁺ neurons. In fact, IL7 is a jugular specific gene highly expressed in both MrgprC11^+^ neurons and MrgprD^+^ neurons. Receptors for IL-4 and IL-7 are distributed across multiple lung immune cell populations, suggesting that cytokines released from jugular sensory neurons could potentially influence diverse immune cells in the lung.

### MrgprC11⁺ and MrgprD⁺ jugular neurons exhibit molecular correspondence with established jugular and DRG nociceptor populations

Previous single-cell transcriptomic studies have identified multiple molecularly distinct nodose and jugular neuron clusters (7, 8, 10, 13). We performed Spearman correlation analysis to determine how the genetically defined jugular neuronal populations isolated in our study correspond to previously identified vagal sensory neuron clusters (Figure 3A). Overall, both MrgprC11⁺ and MrgprD⁺ populations showed consistently higher correlations with jugular than nodose clusters, supporting their molecular identity as jugular sensory neurons. Kupari et al. identified the jugular cluster JG2, characterized by *MrgprD* expression, and JG3, characterized by *Osmr* along with *S1pr1, MrgprA3, Sst,* and *Nppb* (7). Consistently, our MrgprD^+^ group showed the highest correlation score with cluster JG2 (r = 0.75). MrgprC11 was not detected by most of the single-cell transcriptomic analysis including Kupari et al and therefore was not available as a marker for defining their jugular clusters. Our MrgprC11⁺ population did not show a one-to-one correspondence with an individual cluster, but instead showed relatively high correlation scores with JG2, JG3, and JG4 (r = 0.77, 0.79, and 0.71 respectively), highlighting the complementary information provided by genetic and transcriptomic approaches to define sensory neuron populations.

**Figure 3.**
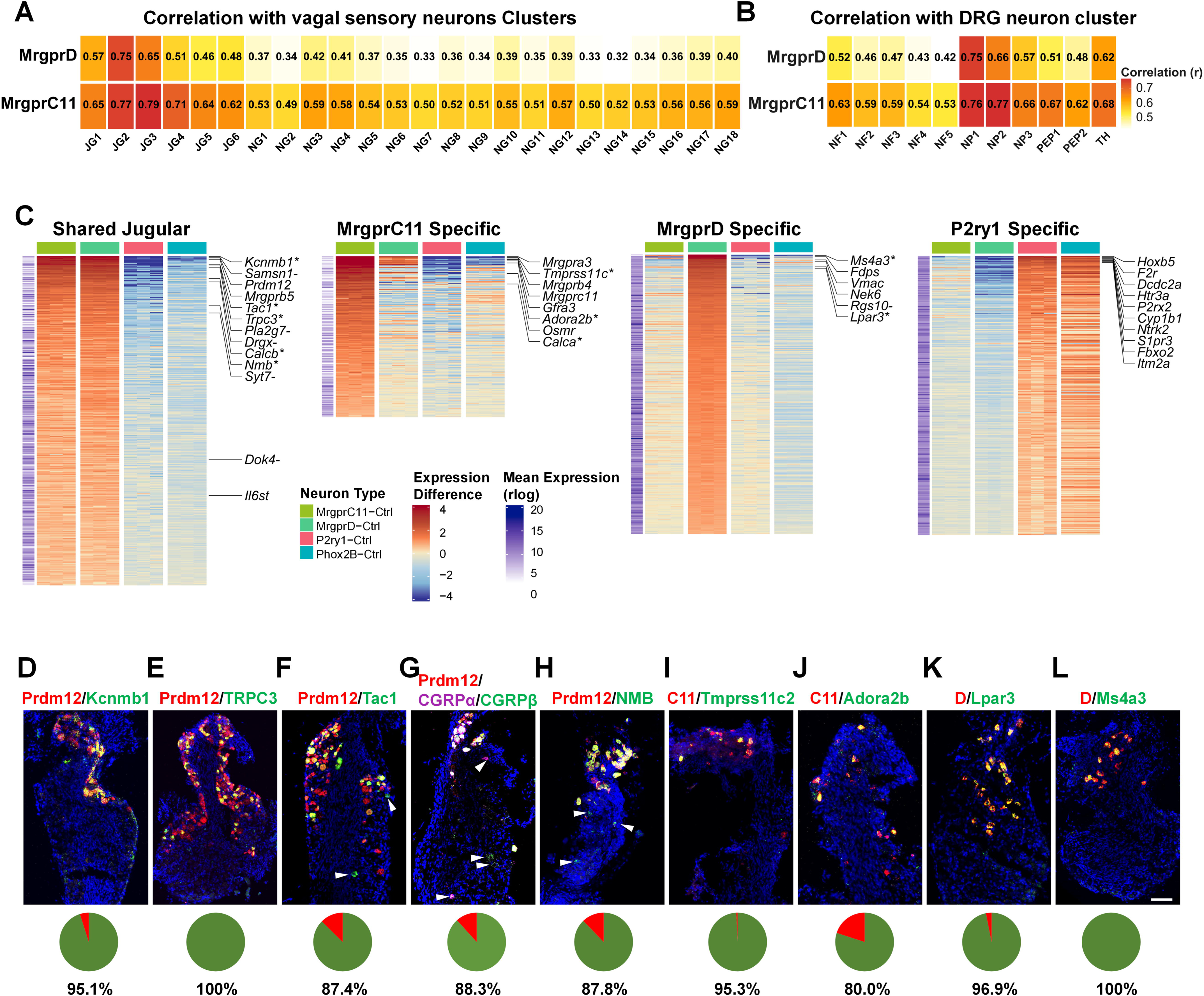
Subtype-Specific Genes (SSGs) signatures across sensory neuron subtypes. (A-B) Correlation heatmaps comparing the transcriptomes of MrgprC11⁺ and MrgprD⁺ jugular neurons with previously defined vagal sensory neuron clusters (A) (Kupari et al., 2019) and DRG neuron clusters (B) (Usoskin et al., 2015). Values represent row-scaled Spearman correlation coefficients (row z-score). (C) Heatmaps showing SSGs. Within each category, genes are ranked according to expression distinctiveness. For each gene, rlog expression values were centered by subtracting the global mean rlog value across all samples, generating a per-sample rlog difference from the mean. Color represents the rlog difference from the mean (red, above mean; blue, below mean). Asterisks (*) denote genes subsequently validated by RNAscope in situ hybridization. Hyphen (-) indicate genes identified by previous studies. Only genes in the top 75% of expressed genes (avg rlog > 3.82) with significant differential expression (padj < 0.05, |log2FC| > 1) are shown. The full gene list for each category is provided in Table S3. (D-L) Top: Representative RNAscope in situ hybridization images of vagal ganglion sections showing the indicated markers. DAPI nuclear staining is shown in blue. In F-H, arrowheads indicate marker-positive cells outside the jugular/Prdm12⁺ cluster. Bottom: Pie charts showing the percentage of cells expressing the indicated marker gene (green) that also co-express the reference gene (red: Prdm12, MrgprC11, or MrgprD). Scale bars, 100 μm.

Jugular and DRG sensory neurons share a neural crest developmental origin, and previous study has demonstrated close correspondence between jugular and DRG neuronal clusters (7, 39). In particular, the JG2/*MrgprD* jugular cluster closely corresponds to the NP1/*MrgprD* DRG population, whereas the JG3/*Osmr/S1pr1/MrgprA3* jugular cluster shows similarity to the NP2/*MrgprA3/MrgprC11* and NP3/*Osmr/S1pr1/MrgprC11/Sst/Nppb* DRG subtypes. Consistent with these relationships, our MrgprD⁺ jugular neurons correlated most strongly with the NP1/*MrgprD* cluster, whereas MrgprC11⁺ neurons showed high correlations with both NP1/*MrgprD* and NP2/*MrgprA3/MrgprC11* DRG clusters (Figure 3B).

### Identification of subtype-specific genes in jugular sensory neurons

Given the limited transcriptomic characterization of jugular sensory neurons to date, we then sought to define the molecular signatures that distinguish the jugular subtypes (MrgprC11^+^ and MrgprD^+^) from nodose neurons (Phox2B^+^ and P2ry1^+^). While the integration of SMART-seq and bulk RNA-seq allowed for the profiling of all the samples, we acknowledge the inherent technical differences in detection sensitivity and library complexity between these methods. To mitigate these effects, we retained highly expressed genes above the 75th percentile and applied regularized log (rlog) transformation (average rlog > 3.82) to stabilize variance across samples with different library sizes. Heatmaps were generated to visualize jugular-enriched gene-expression patterns (Figure 3C, Table S3). Genes were grouped into four categories based on DESeq2 pairwise comparisons: Shared Jugular (enriched in both MrgprC11^+^ and MrgprD^+^ neurons), MrgprC11⁺-specific, MrgprD⁺-specific, and P2ry1⁺-specific genes. All qualifying Shared Jugular and MrgprC11⁺-specific genes were included in the heatmap. Due to the larger size of the MrgprD⁺-specific and P2ry1⁺-specific gene sets, only the top 400 genes ranked by average rlog value were included in the visualization (Figure 3A). The first category consisted of shared jugular-enriched genes, which were similarly enriched in MrgprC11⁺ and MrgprD⁺ neurons compared with the two nodose populations (N = 465 genes; |log2FC| ≤ 1 for MrgprC11⁺ vs. MrgprD⁺). Several genes previously identified by Kupari et al. as jugular-enriched, including Drgx (Prrxl1), Pla2g7, Samsn1, Dok4, Rgs10, and Syt7, were confirmed in our analysis (7). Top-ranked MrgprC11⁺-specific genes (N = 229) included *Mrgprs, Osmr,* and *Calca*, whereas top-ranked MrgprD⁺-specific genes (N = 805) included *Ms4a3, Rgs10,* and *Lpar3*. *Tac1*, previously identified as jugular-enriched and confirmed by our RNAscope analysis (Figure 3F), was manually included in the shared jugular heatmap because variable expression in several nodose samples resulted in Cook’s distance-based outlier flagging and unavailable (NA) adjusted p-values in DESeq2. The *Tac1* raw count and TPM results are provided in Table S3. The P2ry1⁺-specific group (N = 1,430 genes) was defined by comparison with the MrgprC11⁺ and MrgprD⁺ populations only. Phox2b⁺ neurons were excluded from this comparison because P2ry1⁺ neurons represent a subpopulation of Phox2b⁺ nodose neurons. Thus, this group represents genes distinguishing P2ry1⁺ neurons from the two jugular nociceptive populations.

To validate the identified Subtype Specific Genes (SSGs) *in vivo*, we performed RNAscope *in situ* hybridization for a subset of genes selected based on high average rlog expression within each category (Figure 3D-L). Quantification revealed that *Kcnmb1* (95.1%) and *Trpc3* (100%) exhibited nearly complete overlap with Prdm12⁺ neurons, confirming their jugular specificity. *Tac1* (87.4%), *Calca*/*Calcb* (88.3%), and *Nmb* (87.8%) showed 87–89% overlap with *Prdm12*⁺ neurons, indicating that these genes are highly enriched in jugular neurons but are also expressed in a subset of non-jugular neurons. *Tmprss11c* (95.3%) and *Adora2b* (80.0%) were highly enriched in MrgprC11⁺ neurons, whereas *Ms4a3* (100%) and *Lpar3* (96.9%) were specifically expressed in MrgprD⁺ neurons. Notably, many of these SSGs such as *Kcnmb1*, *Tmprss11c*, and *Ms4a3* are identified here as jugular-specific genes for the first time.

We compared our jugular-specific gene sets with genes detected by Kupari et al. in jugular clusters (7). Among the MrgprC11-specific and MrgprD-specific genes identified in our study, 43.5% (450/1,034) were not detected in the previous single-cell sequencing dataset, demonstrating that our high-depth transcriptomic profiling of genetically defined populations provides complementary, higher-sensitivity molecular characterization and enables the detection of low-to moderately expressed genes.

To characterize the functional diversity of jugular-specific genes, SSGs were examined across six curated functional categories, including transcription factors, cell adhesion molecules, GPCRs, ion channels, synaptic vesicle exocytosis genes, and inflammatory response genes (Figure 4, Table S4 and Table S5). Jugular sensory neurons showed distinct expression patterns from nodose neurons across all functional categories. Moreover, MrgprC11⁺ and MrgprD⁺ neurons exhibited substantial molecular differences across these categories, supporting their identities as distinct jugular nociceptive populations with different peripheral innervation patterns and specialized functions.

**Figure 4.**
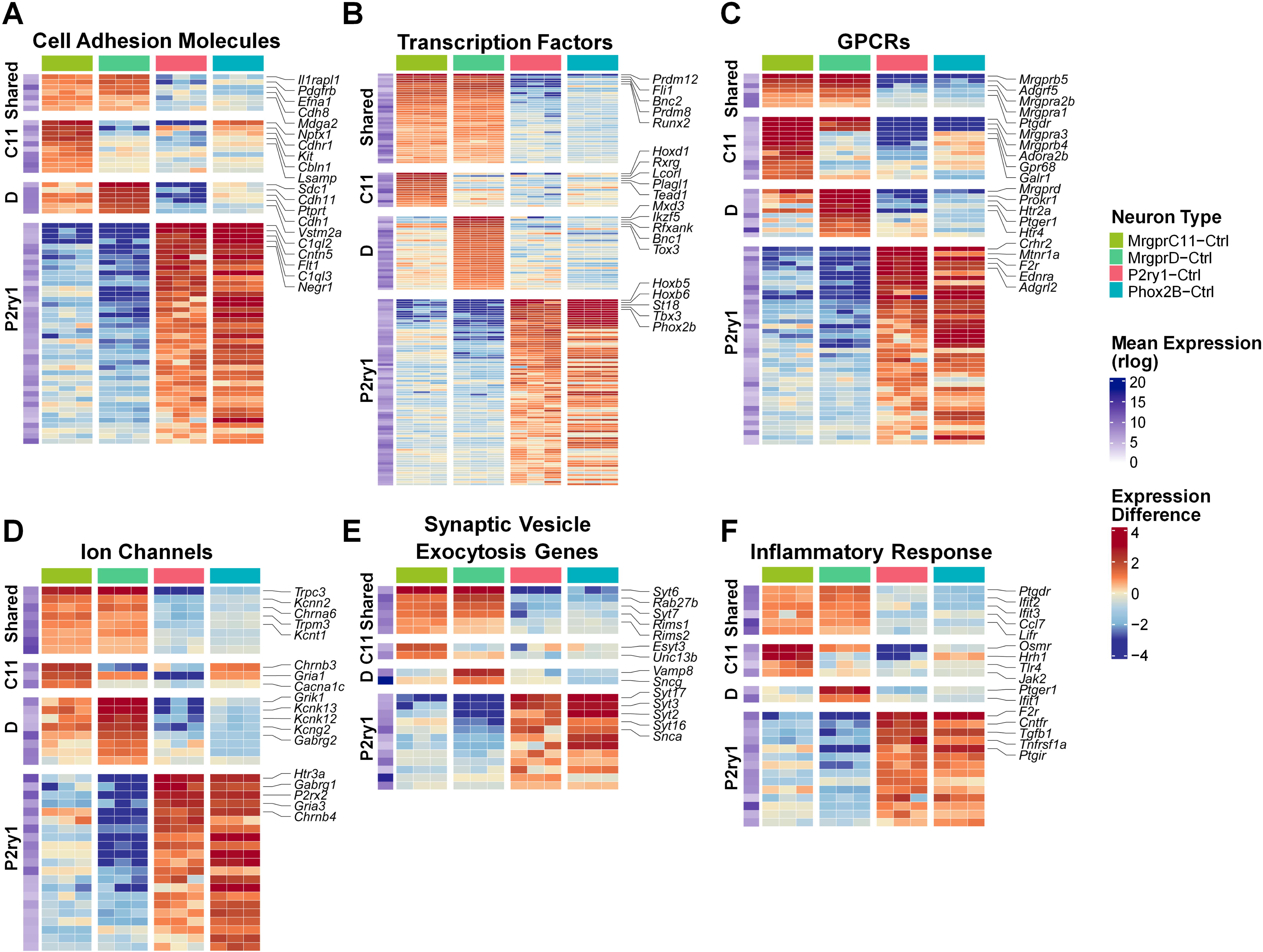
Subtype-specific genes within curated functional categories. Heatmaps showing rlog expression differences from the global mean for SSGs within each curated functional category. Only genes among the top 75% of expressed genes (average rlog > 3.82) with significant differential expression (padj < 0.05, |log2FC| > 1) are shown. The left sidebar indicates mean rlog expression across all four subtypes. The full list of SSGs in each functional category is provided in Table S5.

### MrgprC11⁺ and P2ry1⁺ neurons exhibit divergent transcriptional responses to allergic airway inflammation

To understand the transcriptomic changes of airway-innervating vagal sensory neurons in response to allergic lung inflammation, we analyzed DEGs from naïve versus OVA-treated MrgprC11⁺ and P2ry1⁺ neurons. We found that MrgprC11⁺ and P2ry1⁺ neurons exhibit strikingly divergent transcriptional responses to the same allergic inflammatory challenge (Figure 5). A total of 1797 and 1123 DEGs were identified in MrgprC11^+^ neurons and P2ry1⁺ neurons, respectively. Although a large number of DEGs (431 genes) were shared in both neuronal subtypes, the majority of shared DEGs (418/431, 97%) were regulated in opposite directions, whereas only 13 genes (3%) changed in the same direction (Figure 5A). DEGs from each comparison were divided into four groups (MrgprC11-OVA Upregulated, MrgprC11-OVA Downregulated, P2ry1-OVA Upregulated, and P2ry1-OVA Downregulated) for Gene Ontology Enrichment analysis. The top 20 enriched GO terms for each group were visualized across the Biological Process (BP), Molecular Function (MF), and Cellular Component (CC) ontologies (Figure 5B-D, Table S6). Although many GO terms were enriched in both neuronal populations, they were almost invariably associated with opposite directions of regulation. No GO terms were shared between the upregulated gene sets of the two neuronal populations across the three GO ontologies, and only a single Cellular Component term was shared between the downregulated gene sets.

**Figure 5.**
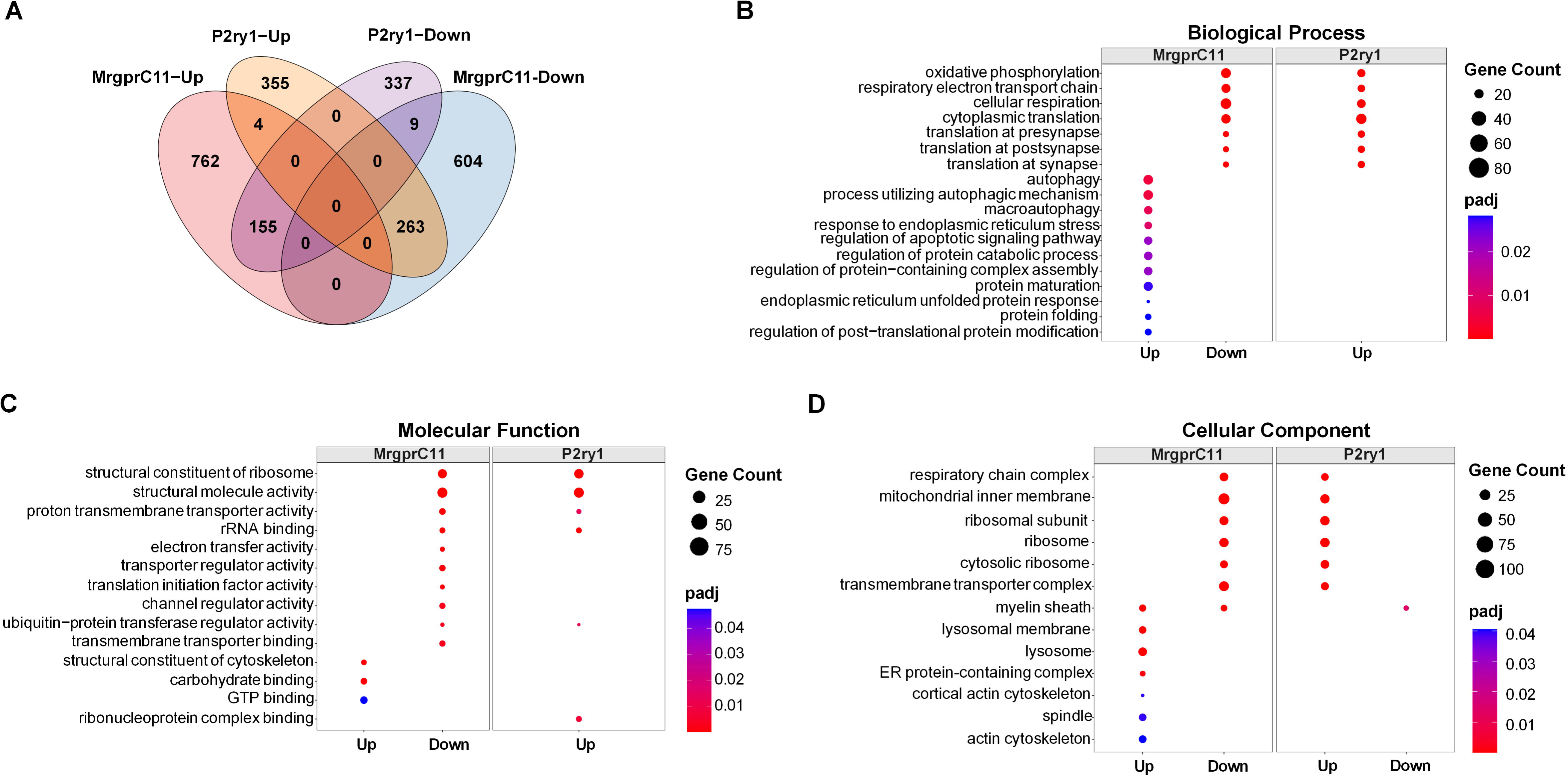
Gene Ontology (GO) enrichment analysis shows distinct subtype-specific transcriptional responses to allergic airway inflammation. (A) Four-way Venn diagram showing the overlap of differentially expressed genes (DEGs). Numbers indicate gene counts unique to or shared among the four DEG sets (MrgprC11-Up, MrgprC11-Down, P2ry1-Up, and P2ry1-Down). (B-D) GO enrichment dot plots comparing upregulated and downregulated gene sets in MrgprC11⁺ and P2ry1⁺ neurons under allergic airway inflammation condition, across three GO categories: (B) Biological Process, (C) Molecular Function, and (D) Cellular Component. Each dot represents a significantly enriched GO term (padj < 0.05). Dot size indicates the number of genes associated with each term; dot color indicates the adjusted p-value. The full GO term list is provided in Table S6.

In MrgprC11⁺ jugular C-fibers, downregulated GO terms were consistently associated with suppression of translational and bioenergetic processes across all three GO ontologies. These terms indicate coordinated suppression of protein synthesis and mitochondrial energy metabolism (40–42). Upregulated GO terms in MrgprC11⁺ jugular C-fibers were strongly associated with activation of protein quality-control mechanisms and degradative pathways involved in the clearance of damaged cellular components (43–45). These results suggest that MrgprC11⁺ jugular neurons adopt a metabolically restrained state, suppressing translation and bioenergetic activity while prioritizing autophagic and proteostatic pathways that may support cellular survival. In contrast, P2ry1⁺ nodose A-fibers exhibited a strikingly opposite response. GO enrichment results of P2ry1⁺ neurons were dominated by upregulated terms, with a few downregulated terms identified. The upregulated GO terms were primarily associated with enhanced translational and bioenergetic processes, suggesting that P2ry1⁺ neurons adopt a metabolically active adaptive program during OVA-induced inflammation (15, 42, 44, 46).

The GO term myelin sheath was enriched in both the upregulated and downregulated Cellular Component (CC) categories of MrgprC11⁺ neurons. Notably, the two MrgprC11⁺ myelin sheath clusters were driven by entirely distinct, non-overlapping gene sets. Furthermore, the downregulated myelin sheath genes in P2ry1⁺ neurons largely overlapped with the upregulated myelin sheath genes in MrgprC11.

Together, these findings indicate that allergic lung inflammation engages similar biological pathways in jugular and nodose airway afferents, but regulates them in largely opposite directions, resulting in fundamentally distinct subtype-specific transcriptional responses.

### Allergic inflammation-induced transcriptomic changes in functional categories

To further characterize OVA-responsive genes, we examined DEG distributions across 7 functional categories and visualized expression changes using subtype-paired heatmaps (Figure 6, Table S7). Across all categories, MrgprC11⁺ neurons consistently showed more DEGs than P2ry1⁺ neurons, and most shared DEGs were regulated in opposite directions, marked by † in the heatmaps. The median absolute *log*_2_*Fold C*ℎ*ange* was 0.41 for MrgprC11 and 0.78 for P2ry1. This modest shift suggests that allergic inflammation fine-tunes existing transcriptional programs while preserving the molecular identities of these vagal sensory neuron subtypes (39, 47).

**Figure 6.**
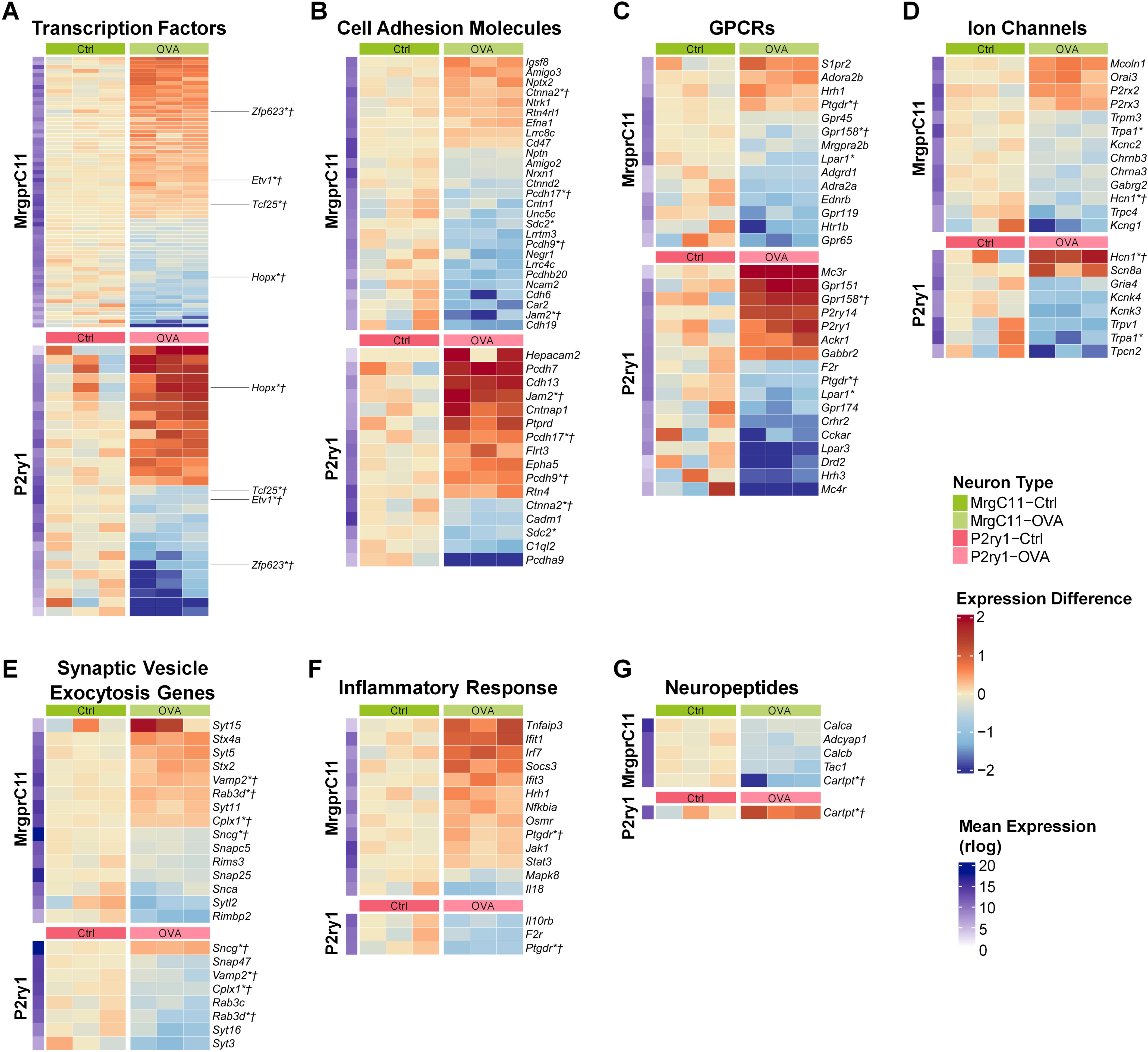
OVA-induced differential gene expression across functional categories in MrgprC11⁺ and P2ry1⁺ airway afferents. (A-G) Heatmaps showing OVA-induced differentially expressed genes (DEGs) in MrgprC11⁺ and P2ry1⁺ neurons across seven curated functional categories. Within each category, genes are ranked by expression difference between the two neuron populations. The left sidebar indicates mean rlog expression level. Only genes in the top 75% of expressed genes (avg rlog > 2.52) with significant differential expression (padj < 0.05) are shown. Genes significant in both populations were marked (*), and those regulated in opposite directions between the two populations were additionally flagged (†). Full list of DEGs in all functional categories is provided in Table S7.

Within these functional categories, many OVA-responsive genes are involved in detecting chemical irritants and inflammatory signaling. In MrgprC11⁺ neurons, OVA increased the expression of *S1pr2* (sphingosine-1-phosphate receptor 2)*, Adora2b* (adenosine A2B receptor)*, Hrh1* (histamine H1 receptor), and *Ptgdr* (prostaglandin D2 Receptor, DP1) in the GPCR category and *Osmr* (oncostatin M receptor) in the inflammatory response category, suggesting enhanced responsiveness to inflammatory mediators following allergen challenge. Changes were also observed in molecules involved in neurogenic inflammation. *Tac1 (tachykinin precursor 1), Calca,* and *Calcb* (calcitonin-related polypeptide α and β) encode neuropeptides substance P and CGRP, which are highly enriched in jugular neurons, particularly MrgprC11⁺ neurons. All three genes were downregulated in MrgprC11⁺ neurons. Interestingly, *Trpa1* (transient receptor potential ankyrin 1), a key mediator of chemical irritants and neurogenic airway hypersensitivity (48), was downregulated in both MrgprC11⁺ jugular and P2ry1⁺ nodose neurons following OVA treatment, suggesting a compensatory response to persistent inflammatory stimulation that limits excessive TRPA1-mediated sensory activation.

Beyond these changes associated with chemosensing, many genes exhibited opposing regulation between MrgprC11⁺ and P2ry1⁺ neurons. *Ptgdr* was regulated in opposite directions in MrgprC11⁺ and P2ry1⁺ afferents, suggesting different regulation of PGD2 signaling in the two populations. *Cartpt* (cocaine-and amphetamine-regulated transcript CART peptide), which is highly enriched in P2ry1⁺ neurons compared with the MrgprC11⁺ and MrgprD⁺ neurons, was the only neuropeptide gene altered by airway inflammation in P2ry1⁺ neurons. Interestingly, OVA further increased *Cartpt* expression in P2ry1⁺ neurons while decreasing its expression in MrgprC11⁺ neurons. Similarly, *Hcn1* (hyperpolarization-activated cyclic nucleotide-gated channel 1), which contributes to the hyperpolarization-activated inward current that regulates resting membrane potential and neuronal excitability (49) was downregulated in MrgprC11⁺ neurons but upregulated in P2ry1⁺ neurons, suggesting that allergic inflammation differentially alters the electrical properties of the two airway afferent populations. Together, these subtype-specific changes further illustrate the distinct molecular adaptations of jugular and nodose airway afferents to the same inflammatory challenge.

## Discussion

Because jugular sensory neurons are relatively sparse and technically challenging to isolate, their molecular profiles remain poorly defined. Here, by combining genetic labeling, FACS isolation, and SMART-seq, we performed deep transcriptomic profiling of MrgprC11⁺ and MrgprD⁺ jugular nociceptors. Our results revealed unique subtype-specific molecular signatures. Compared with Phox2b⁺ nodose neurons and P2ry1⁺ airway-innervating nodose A-fibers, both jugular populations exhibited distinct expression patterns across diverse functional categories. Moreover, MrgprC11⁺ and MrgprD⁺ neurons were also molecularly distinct from each other, supporting their identities as different jugular subtypes with specialized axonal innervation patterns and sensory functions.

An interesting feature of MrgprC11⁺ jugular neurons is their similarity to MrgprC11⁺ DRG neurons. Our transcriptomic analysis revealed a shared molecular profile enriched in multiple receptors and ion channels involved in nociceptive and inflammatory sensing. The two populations also exhibit similar peripheral termination patterns. MrgprC11⁺ DRG neurons form free nerve endings in the superficial epidermis, whereas MrgprC11⁺ jugular neurons form free nerve endings along the airway epithelium, with terminals extending toward the luminal surface where they are positioned to detect airway irritants. Despite innervating different organs, both populations mediate nocifensive responses: activation of MrgprC11⁺ nerves in the skin evokes itch and scratching, whereas activation of MrgprC11⁺ jugular nerves in the airway induces bronchoconstriction. Together, these similarities suggest that jugular neurons may represent a visceral extension of neural crest-derived somatosensory nociceptor programs. MrgprC11⁺ DRG and jugular neurons share a transcriptomic program for detecting nociceptive signals, while engaging distinct neural circuits to evoke behavioral or physiological responses specific to their target organs.

MrgprD is expressed in approximately 37% of jugular neurons, representing a large population of jugular C-fibers. Our study provides a detailed transcriptomic characterization of MrgprD⁺ jugular neurons and reveals a distinct molecular profile equipped with key molecules involved in nociceptive sensory functions. However, the physiological functions of MrgprD⁺ jugular neurons remain unknown, and even their peripheral targets have not been identified. A previous study examined the axonal projections of MrgprD⁺ sensory neurons using the *MrgprD^EGFPf^* mouse line and found MrgprD⁺ nerve endings in the epidermis, with a complete absence of innervation in other tissues examined, including the cornea, meninges, stomach, intestine, muscle, lung, heart, and bladder. These findings suggest that MrgprD⁺ jugular neurons may have highly restricted peripheral territories that remain unidentified. Further investigation of the peripheral targets and physiological functions of MrgprD⁺ jugular neurons will help define the unique roles of jugular sensory neurons.

Our results reveal strikingly divergent adaptive responses to allergic airway inflammation between MrgprC11⁺ and P2ry1⁺ neurons. At the pathway level, GO enrichment analysis suggested that MrgprC11⁺ neurons shifted toward a metabolically restrained state, with reduced translational and bioenergetic activities and increased autophagic and proteostatic pathways. In contrast, P2ry1⁺ neurons exhibited a more metabolically active state, with enhanced translation and bioenergetic activities. This divergence was even more striking at the gene level: among 431 genes differentially expressed in both populations following OVA challenge, 418 (97%) were regulated in opposite directions. These genes spanned multiple functional categories, indicating broad subtype-specific transcriptional remodeling rather than differences restricted to individual pathways. Together, these analyses suggest that airway-innervating jugular nociceptors (MrgprC11⁺) and nodose A-fibers (P2ry1⁺) engage largely distinct and opposing transcriptional responses during allergic airway inflammation. This divergence may reflect their intrinsic differences in developmental origin, physiological properties, and sensory functions.

The high expression of proinflammatory neuropeptides, cytokines, and cytokine receptors in vagal sensory neurons highlight the bidirectional communication between sensory nerves and the inflamed lung. Sensory nerves function not only as sensors of irritants and inflammatory signals but also as active participants in shaping the pulmonary immune environment. For example, sensory neuron-derived CGRP can suppress ILC2 activation and type 2 airway inflammation, whereas substance P promotes allergen-induced goblet cell hyperplasia and mucus production (50–56). Both CGRP and substance P are highly enriched in jugular neurons, particularly MrgprC11⁺ neurons, highlighting the important role of jugular nociceptors in regulating airway inflammation. In addition, distinct repertoires of cytokines and cytokine receptors were observed in MrgprC11⁺ and P2ry1⁺ neurons, consistent with their strikingly divergent transcriptomic responses to allergic airway inflammation.

In summary, our transcriptomic study provides a comprehensive molecular characterization of genetically defined jugular nociceptor populations and reveals their distinct molecular signatures.

Moreover, the strikingly divergent transcriptional responses of MrgprC11⁺ jugular and P2ry1⁺ nodose airway afferents to allergic inflammation demonstrate that distinct vagal sensory neuron subtypes adapt differently to the same inflammatory environment. Together, these findings advance our understanding of vagal sensory neuron diversity and specialization in airway sensation and inflammatory responses.

## MATERIALS AND METHODS

### Mice

All experiments were approved by the Georgia Institute of Technology Animal Use and Care Committee. Mice were housed in a vivarium with a 12-h light/dark cycle, in groups of up to five per cage, with food and water available *ad libitum*. All experiments were conducted using 2-6 months old (20-30g) mice. *MrgprC11^CreERT2^* BAC transgenic line was generated by a previous study (57). Mice listed below were purchased from Jackson Laboratory: wild type C57BL/6 mice (Stock No: 000664), *MrgprD^CreERT2^* (Stock No: 031286) (30), *Phox2B^Cre^* (Stock No: 016223), *P2ry1^Cre^* (Stock No: 029284), *ROSA26^tdTomato^*(Stock No: 007909). All transgenic mice used for these experiments have been backcrossed to C57BL/6 mice for at least 10 generations.

### Tamoxifen treatments

To label MrgprC11^+^ neurons and MrgprD^+^ neurons, the *MrgprC11^CreER^*or *MrgprD^CreER^* mice were treated daily via oral gavage (22g x 25mm, FTP-22-25, Instech Laboratories) with tamoxifen (T5648, Sigma) dissolved in sunflower seed oil (S5007, Sigma) for 5 days at 3-4 weeks of age. Tamoxifen concentration used was 100 mg/kg for *MrgprC11^CreER^* and 40 mg/kg for *MrgprD^CreER^*mice.

### Ovalbumin (OVA)-induced allergic lung inflammation

Mice were sensitized to ovalbumin (OVA) by i.p. injection of 50 μg OVA plus 2 mg Imject Alum in 150 μl PBS on Days 1, 7, and 14. Lightly anesthetized mice (isoflurane) were challenged by intratracheal instillation of 200 μg OVA in 50 μl PBS on Days 21, 22, and 23. Tissues were collected after the final challenge for downstream analysis.

### RNAscope *in situ* hybridization

RNAscope *in situ* hybridization was performed using the RNAscope® Multiplex Fluorescent Reagent V2 (ACD 323270) according to the manufacturer’s instructions. Briefly, vagal ganglia were quickly collected and freshly frozen in OCT medium. The ganglia were sectioned at a thickness of 20 µm and the following probes were used: Prdmd12 (524371), MrgprC11 (488771), MrgprD (417921), P2ry1 (406061), TRPV1 (313331), Kcnmb1 (1138061), TRPC3 (525201), Tac1 (410351), CGRPα (578771), CGRPβ (425511), NMB (459931), Tmprss11c2 (1222011), Adora2b (445281), Lpar3 (432591), Ms4a3 (438181). Fluorescent images were acquired on a Confocal Microscopy Zeiss LSM700 and analyzed in Fiji. DAPI staining was performed to visualize the nuclei of the cells. Cells with ≥ 5 fluorescent dots are considered positive. 3000-4000 vagal sensory neurons (3-6 vagal ganglia) were quantified to examine the percentage of neurons expressing each gene. Only cells containing nuclei labeled by DAPI were counted to avoid repetitive counting of the same cells in different sections.

### Fluorescence-activated cell sorting (FACS) and sequencing strategy

For tissue collection, vagal ganglia were freshly dissected and treated with dissociation enzyme solution (5 mg/ml Dispase, 1 mg/ml Collagenase Type I in HPBS without Ca^2+^ and Mg^2+^, Gibco) at 37°C for 40 minutes under constant rotation. The digestant was centrifuged at 200 g for 5 minutes and the precipitant was carefully resuspended in culture medium (90% DMEM/F-12, 10% FBS) before passing through a 100 μm cell strainer. Sensory neurons were subsequently enriched using Percoll gradient centrifugation (30% Percoll). Briefly, the cell suspension (3 mL) was carefully layered onto the Percoll solution (3 ml) to maintain a clear interface and centrifuged at 700 × g for 10 minutes at 4°C with low acceleration and deceleration settings. After centrifugation, non-neuronal cells and fiber-enriched material remained in the upper layer, while cell bodies of vagal sensory neurons formed a pellet at the bottom of the tube. The neuronal pellet was collected, washed by centrifugation, and resuspended in cell sorting buffer. Samples were kept on ice and immediately transported to the flow cytometry facility. Fluorescence-activated cell sorting (FACS) was performed using a BD FACSMelody cell sorter with a 100 μm nozzle. Propidium iodide (PI) staining was used to exclude non-viable cells.

To isolate P2ry1⁺ airway-innervating neurons, 50 μL of CTB-488 (2 μg/mL) was delivered via intratracheal instillation into *P2ry1^tdTomato^* mice one week before tissue collection to retrogradely label airway-innervating vagal sensory neurons. Vagal ganglia from two mice were pooled for each biological replicate and tdTomato⁺/CTB-488⁺ neurons were isolated by FACS. Due to the low abundance of MrgprC11⁺ and MrgprD⁺ vagal sensory neurons, selective isolation of airway-innervating subsets was not feasible. Instead, all tdTomato⁺ neurons were collected from *MrgprC11^tdTomato^* and *MrgprD^tdTomato^* mice. Vagal ganglia from 5-6 mice were pooled for each biological replicate, yielding approximately 90-170 sorted cells per sample. Three biological replicates were collected for each neuronal population. Cells were sorted directly into lysis buffer (Takara, 635013) and shipped to Admera Health for library preparation and SMART-seq (35). cDNAs were synthesized using the SMART-Seq v4 Ultra Low Input RNA Kit (Takara) which enables high-quality cDNA synthesis directly from sorted cells without RNA extraction.

Phox2B⁺ neurons were collected for sequencing to characterize the transcriptome of nodose neurons and to establish a reference for comparison with jugular MrgprC11⁺ and MrgprD⁺ neurons. Because the nodose ganglion comprises thousands of neurons representing many molecularly distinct subtypes, a large number of Phox2B⁺ neurons were collected to maximize coverage of the endogenous nodose neuronal diversity. Vagal ganglia from two *Phox2B^tdTomato^* mice were pooled per sample, yielding approximately 4,000 tdTomato⁺ cells per sample. Because SMART-seq is not suitable for samples containing more than 1,000 cells, the Phox2B⁺ population was profiled by bulk RNA sequencing rather than SMART-seq. RNA was extracted using the Qiagen RNeasy Micro Kit and submitted to Admera Health for library preparation and RNA sequencing.

While MrgprD^+^ and Phox2B^+^ populations were collected under naïve conditions, MrgprC11^+^ and P2ry1^+^ subtypes were isolated from both naïve and ovalbumin (OVA)-sensitized mice. All experimental conditions were analyzed using three independent biological replicates (n = 3), a total of 18 samples. 150-cycle paired-end sequencing was carried out on an Illumina HiSeq 2500 platform, generating approximately 40 million paired-end reads per sample.

### RNA sequencing and data processing

Sequence quality was evaluated using FastQC (version 0.11.8) to identify potential technical artifacts. Trimmomatic (version v0.38) was applied to cut adaptors and trim low-quality bases with default setting (58). STAR Aligner (version 2.7.1a) was used to align the reads. Read summarization was performed using HTSeq-count (version 0.11.2) (59). Raw read count matrices were normalized downstream for unified comparative analysis. Gene expression abundances were mathematically converted to Transcripts Per Million (TPM) values. Sequencing data quality was assessed by examining the total mapped reads per sample (average=32,324,309; range: 17,824,244-46,028,043). Genes with mean read count >10 and mean TPM > 0.1 were retained for downstream analysis, yielding 19779 genes. Gene detection was consistent across all samples: MrgprC11^+^ (mean 20,805 genes, range 20,457-21,003), P2ry1^+^ (mean 18,700 genes, range 17,204-20,193;), MrgprD^+^ (mean 20,345 genes, range 19,994-20,742), and Phox2B^+^ (mean 23,374 genes, range 22,837-24,046).

All statistical analyses were performed in R (version 4.6.0) using Bioconductor packages DESeq2 (version 1.52.0), clusterProfiler (version 4.20.0), and org.Mm.eg.db (version 3.23.0) for gene annotation (60, 61). Sample-level quality assessment, including principal component analysis (PCA) and hierarchical clustering of sample correlations and distances, was performed using variance-stabilizing transformation (VST) implemented in DESeq2. Differential expression analysis was conducted using DESeq2. Baseline comparisons among the four neuronal subtypes (MrgprC11, MrgprD, P2ry1, and Phox2B) were performed using DESeq2 with a single joint model fit across all 12 samples with pairwise contrasts extracted between each subtype pair from this joint model.

Raw read counts from all samples were combined into a single DESeq2 object and transformed using the regularized logarithm (rlog) with blind dispersion estimation to stabilize variance across the expression range (68). Only genes within the top 75th percentile of the average rlog distribution (average rlog > 3.82) were considered for enrichment analysis, yielding a candidate set of 14,317 genes. For each gene, an enrichment score was calculated as the difference between the individual sample rlog and the global mean rlog across all subtypes (*rlog_diff_* = *rlog_i_* − *rlog_mean_*). Genes were classified into four categories based on DESeq2 pairwise comparisons with an adjusted p-value threshold of *padj* < 0.05 and a |*log*_2_*Fold C*ℎ*ange*| > 1: (1) Shared Jugular genes were significantly enriched in both MrgprC11 and MrgprD relative to both P2ry1 and Phox2B, with similar expression between the two jugular subtypes (|*log*_2_*Fold C*ℎ*ange*| ≤ 1); (2) MrgprC11-specific genes were significantly enriched in MrgprC11 relative to P2ry1, Phox2B, and MrgprD; and (3) MrgprD-specific genes were significantly enriched in MrgprD relative to P2ry1, Phox2B, and MrgprC11; and (4) P2ry1-Specific genes were significantly enriched in P2ry1 relative to MrgC11 and MrgD. Phox2b⁺ neurons were excluded from this comparison because P2ry1⁺ neurons represent a subpopulation of Phox2b⁺ nodose neurons.

Heatmaps were generated using pheatmap (version 1.0.13) and complexHeatmap in R (version 2.28.0), and additional visualization was performed using ggplot2 (version 4.0.3) (35, 61, 62). Heatmaps displayed the rlog difference from the mean, with values clipped to a scale of −3 to 3 to emphasize relative enrichment. To provide context for absolute expression magnitude, heatmaps included sidebar annotations for the Mean Expression (rlog). Genes were ordered by descending specificity score within each category. All genes classified as Shared Jugular or MrgprC11⁺-specific were included in the heatmap visualization. Because the MrgprD⁺-specific and P2ry1⁺-specific categories contained substantially larger gene sets, both were restricted to the top 400 genes ranked by mean rlog expression across all samples prior to plotting. Within each category, selected genes were ranked using a specificity sort score reflecting relative enrichment in the target population versus all other groups, defined as follows:

Shared Jugular: mean rlog (MrgprC11, MrgprD) − mean rlog (P2ry1, Phox2b) MrgprC11⁺-specific: mean rlog (MrgprC11) − mean rlog (MrgprD, P2ry1, Phox2b) MrgprD⁺-specific: mean rlog (MrgprD) − mean rlog (MrgprC11, P2ry1, Phox2b) P2ry1⁺-specific: mean rlog (P2ry1) − mean rlog (MrgprC11, MrgprD) For Shared Jugular genes, the score was calculated as the average rlog difference in MrgprC11⁺ and MrgprD⁺ neurons relative to the nodose populations. For subtype-specific genes, the score was calculated as the rlog difference in the target subtype relative to the average of the remaining three populations. Genes were ordered on the heatmaps by decreasing enrichment score, with the most selectively enriched genes displayed at the top of each category.

### Cross-dataset correlation with published sensory neuron atlases

Pseudobulk expression profiles were generated by averaging single-cell expression within each published cluster from Kupari et al. (2019, vagal ganglion; jugular-derived JG1–6, nodose-derived NG1–18) and Usoskin et al. (2015, DRG; NF1–5, NP1–3, PEP1–2, TH), restricted to genes shared with the jugular bulk dataset (7, 39). Both the jugular bulk TPM values and the published pseudobulk profiles were log2(x+1)-transformed prior to correlation, placing the two independently generated datasets on a comparable scale. Spearman correlation was computed between each jugular population’s (MrgprC11, MrgprD) and each published cluster’s log2(x+1)-transformed profiles; values shown are row-scaled (z-score) for visualization, with the underlying Spearman r annotated in each cell (63).

### Curated functional categories

Cell adhesion molecules and synaptic vesicle exocytosis genes were defined using the curated hippocampal gene-family datasets of Földy et al. (64). G protein-coupled receptors and ion channels were defined using the IUPHAR/BPS Guide to Pharmacology database (version 2026.2) (65). Transcription factors were defined using the mouse sequence-specific DNA-binding transcription factor catalogue from AnimalTFDB 4.0 (66). Neuropeptide genes were manually curated using established nomenclature resources. To establish the inflammatory response category, we integrated genes annotated to cytokine receptor activity (GO:0004896) and inflammatory response (GO:0006954) retrieved from Mouse Genome Informatics (MGI) (67). Each gene was assigned to a single functional category by gene-family membership. The full gene list for all functional categories is provided in Table S5.

### Differential expression analysis of allergic lung inflammation model

Only genes within the top 75th percentile of the average rlog distribution (average rlog > 2.52) were considered for downstream analysis, yielding a candidate set of 16,075 genes. DEGs were defined as those with *padj* < 0.05. No fold-change threshold was applied, as OVA sensitization is expected to produce moderate transcriptional changes within existing neurons rather than the large expression differences observed between distinct cell populations. The resulting expression filter threshold (average rlog > 2.52; 16,075 genes) differs slightly from the jugular-subtype analysis (average rlog > 3.82; 14,317 genes) because the smaller sample set and different condition structure produce a different rlog distribution.

Gene ontology (GO) enrichment analysis was performed using the compareCluster function in the clusterProfiler R package, with separate analyses for upregulated and downregulated gene sets in each comparison. DEGs from each comparison were divided into four groups, MrgprC11-OVA Upregulated, MrgprC11-OVA Downregulated, P2ry1-OVA Upregulated, and P2ry1-OVA Downregulated. Enrichment was tested for Biological Process (BP), Cellular Component (CC), and Molecular Function (MF) ontologies using mouse gene annotations (org.Mm.eg.db). Terms were considered significant at adjusted p-value threshold of *padj* < 0.05, with gene set sizes restricted to 3–500 to exclude overly specific or overly broad terms (68). The top 20 most significant terms per group were selected for visualization, and redundant or non-neuronal terms were manually curated to improve interpretability. Dot plots showing GO analyses were generated with ggplot2, with dot size representing gene count and color indicating adjusted p-value.

## Conflicts of interest

The authors claim no conflict of interest.

## Funding

The work was supported by grants from the US National Institutes of Health (HL173002) and National Science Foundation (2334697) to L.H.

## Acknowledgments

We thank the Department of Animal Resources at the Georgia Institute of Technology for the animal care and services.

## Notes

### Competing Interest Statement

The authors have declared no competing interest.

